# MetaGate: Interactive Analysis of High-Dimensional Cytometry Data with Meta Data Integration

**DOI:** 10.1101/2023.10.27.564454

**Authors:** Eivind Heggernes Ask, Astrid Tschan-Plessl, Hanna Julie Hoel, Arne Kolstad, Harald Holte, Karl-Johan Malmberg

## Abstract

Flow cytometry is a powerful technology for high-throughput protein quantification at the single-cell level, widely used in basic research and routine clinical diagnostics. Traditionally, data analysis is carried out using manual gating, in which cut-offs are defined manually for each marker. Recent technical advances, including the introduction of mass cytometry, have increased the number of proteins that can be simultaneously assessed in each cell. To tackle the resulting escalation in data complexity, numerous new analysis algorithms have been developed. However, many of these show limitations in terms of providing statistical testing, data sharing, cross-experiment comparability integration with clinical data. We developed MetaGate as a platform for interactive statistical analysis and visualization of manually gated high-dimensional cytometry data with integration of clinical meta data. MetaGate allows manual gating to take place in traditional cytometry analysis software, while providing a combinatorial gating system for simple and transparent definition of biologically relevant cell populations. We demonstrate the utility of MetaGate through a comprehensive analysis of peripheral blood immune cells from 28 patients with diffuse large B-cell lymphoma (DLBCL) along with 17 age- and sex-matched healthy controls using two mass cytometry panels made of a total of 55 phenotypic markers. In a two-step process, raw data from 143 FCS files is first condensed through a data reduction algorithm and combined with information from manual gates, user-defined cellular populations and clinical meta data. This results in one single small project file containing all relevant information to allow rapid statistical calculation and visualization of any desired comparison, including box plots, heatmaps and volcano plots. Our detailed characterization of the peripheral blood immune cell repertoire in patients with DLBCL corroborate previous reports showing expansion of monocytic myeloid-derived suppressor cells, as well as an inverse correlation between NK cell numbers and disease progression.

## Introduction

Fluorescence-based flow cytometry was invented in the late 1960s, and has since gained widespread popularity in basic research, routine diagnostics and clinical trials. Modern flow cytometers allow simultaneous quantification of more than 40 antigens with single-cell resolution, and the introduction of mass cytometry has further increased this number.^1^ This has enabled detailed functional and phenotypic characterization of very complex subsets of cells within highly heterogenous sample material, such as peripheral blood or tumor tissue.

The massive advances in cytometry technology have posed challenges for bioinformatical analysis. Traditionally, cytometry data analysis is carried out by manually defining biologically relevant cell populations by setting cut-off values for multiple antigen markers. This strategy, termed manual gating, allows consideration of known biology, internal controls, and experiment-specific technical issues in the data analysis. However, with increasing data complexity, manual gating becomes labor-intensive and prone to operator bias.^1–3^ In response to these challenges, a vast collection of clustering and dimensionality reduction algorithms has been implemented for cytometry data analysis and visualization, including *t-SNE*, *PhenoGraph*, *SPADE* and *FlowSOM*.^4–8^ Although representing major advances in our ability to explore and understand high-dimensional single-cell data, the output of these algorithms can be unpredictable, due to experiment-specific marker selection, technical variation or inherent properties of different clustering methods.^9^

Despite its limitations and the plethora of new analysis algorithms available, manual gating remains the most widely used method for cytometry data analysis. However, stratification of samples, statistical analysis and visualization of summarized data typically involves multiple data handling steps in different software packages, potentially reducing throughput and data traceability. To alleviate these problems, we developed the MetaGate R package. Through its graphical user interface, MetaGate provides a platform for statistical analysis and visualization of complex cytometry data sets from raw data via feature selection to publication-ready figures, based on manual gating performed in two of the most popular flow cytometry analysis software packages, FlowJo and Cytobank.

Along with genomics, proteomics and immunological imaging techniques, cytometry remains a crucial tool for assessing the immune system in cancer, both within the tumor microenvironment and at the global level. Such understanding is important for cancer prevention, diagnostics, prognostication and development of novel treatment strategies. To display the capabilities of MetaGate in such studies, we performed a broad mass cytometry characterization of peripheral blood from a cohort of 28 patients with diffuse large B-cell lymphoma (DLBCL) alongside 17 healthy blood donors.

DLBCL is the most common group of non-Hodgkin lymphoma, with an incidence in the United States of around 7 cases per 100,000 persons per year.^10^ First-line treatment usually includes multi-agent chemotherapy in combination with the anti-CD20 monoclonal antibody rituximab. Two main subtypes, germinal-center B-cell (GCB) and activated B-cell (ABC) type, have been identified, correlating fairly well with histological features and explaining some of the outcome variation.^11^ However, the highly diverse presentation and outcome, which cannot fully be explained by existing clinical, histological or biochemical markers, remains a major clinical challenge.^12^ Therefore, to improve diagnostics, prognostics and treatment of this disease, there is a need for a better understanding of the heterogeneity of its presentation and immunological responses.

The mass cytometry data from this study, which in part is previously published,^13^ is analyzed using MetaGate and describes a substantial impact on the immune system from both the disease and its treatment. All data figures and statistical analyses are generated in the MetaGate user interface. The MetaGate R package and source code is made publicly available, along with all mass cytometry data and meta data, enabling anyone to reproduce the analysis, as well as further develop or use MetaGate for other data sets.

## Methods

### Development of MetaGate

MetaGate is developed as an R^14^ package with a web browser-based graphical user interface implemented using the *shiny* package.^15^ Interaction with FlowJo workspaces, GatingML files and Flow Cytometry standard (FCS) files is implemented with the use of the *flowWorkspace*, *CytoML*, *flowCore* and *flowUtils* packages.^16–19^ Plots are generated using the *ggplot2* package.^20^

### Patient samples and clinical data

The use of patient and healthy donor blood samples and clinical data was approved by the regional ethical board in Norway (ref. 2012/1143, 2015/2142, 2018/2482 and 2018/2485). Patients were selected from a lymphoma patient biobank established in January 2015 at Oslo University Hospital. Fully informed written consent was obtained from all healthy donors and patients. The study includes 17 healthy donors and 28 patients. Median age was 65 for healthy donors and 67 for patients, while the percentages of female subjects were 53% and 43%, respectively. Peripheral blood mononuclear cells (PBMC) were collected from patients directly before initiation and after completion of first-line chemotherapy, while healthy donor samples were collected at one timepoint. Inclusion diagnoses were diffuse large B-cell lymphoma (DLBCL), high-grade B-cell lymphoma (HGBCL) with MYC and BCL2 and/or BCL6 rearrangements (or based on the 2008 WHO classification of lymphoid neoplasms, “B-cell lymphoma, unclassifiable, with features intermediate between diffuse large B-cell lymphoma and Burkitt lymphoma”), and T-cell/histiocyte-rich large B-cell lymphoma (THRLBCL). All patients were treated with a combination of rituximab and chemotherapy regimens containing cyclophosphamide, doxorubicin, vincristine, etoposide and prednisolone (CHOP/EPOCH/CHOEP). The Hans algorithm was used for subtype classification of germinal center B-cell like (GCB) and non-GCB DLBCL. For patients, absolute numbers of lymphocytes were retrieved from diagnostic white blood cell differential counts, while such data was not available for healthy donors.

### Mass cytometry

PBMC from patients and healthy blood donors were isolated by density gradient centrifugation using Lymphoprep (Axis-Shield, Oslo, Norway). Cells were subsequently aliquoted and cryopreserved in 10% DMSO, 70% fetal calf serum (FCS) (Sigma-Aldric, St. Louis, MO) and 20% RPMI 1640 (Thermo Fisher Scientific, Waltham, MA). Upon experiments, PBMCs were thawed and rested over-night in RPMI 1640 with 10% FCS.

Cells were stained with Cell-ID Intercalator-Rh (Fluidigm, San Francisco, CA) and GLUT1.RBD.GFP (Metafora Biosystems, Evry cedex, France) according to the manufacturer’s instructions to allow for viability testing and GLUT-1 detection, respectively. Samples were then incubated with an Fc receptor binding inhibitor polyclonal antibody (Thermo Fisher Scientific), before staining with a surface antibody cocktail (Supplementary Table 1). Antibodies were either obtained pre-labeled from Fluidigm or in-house conjugated using Maxpar X8 antibody labeling kits (Fluidigm). After staining, the cells were fixed using 2% paraformaldehyde in PBS without Ca and Mg), and then permeabilized and barcoded using the Cell-ID 20-Plex Barcoding Kit (Fluidigm) according to the manufacturer’s instructions. Samples were then pooled, resuspended in pure methanol and stored at −20°C. On the day of mass cytometry acquisition, samples were thawed, stained with an intracellular antibody cocktail and labeled with Cell-ID Intercalator-Ir (Fluidigm) according to manufacturer’s instructions. Immediately before acquisition, samples were supplemented with EQ Four Element Calibration Beads (Fludigim) and acquired on a CyTOF 2 (Fluidigm), equipped with a SuperSampler (Victorian Airship, Alamo, CA). The event rate was kept below 400 events per second. Samples were analyzed in 8 batches with healthy donors and patients distributed evenly across batches, and patient samples from different timepoints always included in the same batch. Due to lack of sufficient cell numbers, PBMCs from 3 of the healthy donors were not analyzed using mass cytometry panel 2.

### Data preparation

FCS files were normalized using the Fluidigm Helios software, and debarcoded either by manual gating or using the Helios software. The files were then imported in Cytobank (Cytobank, Santa Clara, CA), where debris, doublets and dead cells were excluded. Data was then gated on CD45^+^ events and exported as FCS files. Files from the two panels were imported into separate FlowJo workspaces and gated according to Supplementary Figures 1–2. In each FlowJo workspace, all samples shared identical gating hierarchies, but gates were adjusted manually for each sample. Each FlowJo workspace was then imported in MetaGate. In MetaGate, populations were defined according to Supplementary Tables 3–4. Channels that were empty or representing intercalators or non-relevant markers were excluded (Supplementary Tables 1–2). Furthermore, the markers GLUT-1, CD71, CD137 and NKG2D were removed due to problematic performance or batch effects. Event limit was kept at 50, meaning that populations with less than 50 events were excluded from calculation of marker intensities or child population sizes. No data transformation was applied in MetaGate. Gating strategy plots were generated using the *CytoML* and *ggcyto* R packages.

### Statistical analysis

All statistical plots and statistical analyses were generated in MetaGate version 1.0 on macOS 13.1 running R version 4.2.2. Minor typographical changes and insertion of p value annotation were subsequently performed in Adobe Illustrator version 27.2. The Mann–Whitney *U* test was used for unpaired comparison of two groups (Figures 3B–G, 5A–F). Paired two-group comparisons were tested using the Wilcoxon signed-rank test (Figures 4B–D, 4F). Comparison of multiple groups was done using the Kruskal–Wallis *H* test, and in the case of p values ≤ 0.05 subsequent pairwise group comparisons using the Dunn test (Figure 4A, 4G). Adjustment of p values was not performed.

P values above 0.05 were defined as not significant (ns.), while *, **, *** and **** were used to indicate p values below or equal to 0.05, 0.01, 0.001 and 0.0001, respectively. Bar plot height represents the median, while error bars indicate the inter-quartile range. In box plots, hinges correspond to the 25th and 75th percentile, while whiskers range to the most extreme values, but no longer than 1.5 times the inter-quartile range, and data points outside that range were plotted individually.

### Availability of data and code

The full MetaGate source code is published at https://github.com/malmberglab/metagate. Documentation and installation instructions are available at https://metagate.malmberglab.com. Raw data for the included data set is available from FlowRepository using accession code FR-FCM-Z6DF. The MetaGate file used to generate all statistics and figures can be downloaded from https://metagate.malmberglab.com.

## Results

### Generating a MetaGate data set

MetaGate is based on manual gating, which can be performed in either the FlowJo or Cytobank software packages. Blood samples or other cell suspensions are analyzed using a mass or flow cytometer (Figure 1A), which generates Flow Cytometry Standard (FCS) files. These are imported in FlowJo or Cytobank. After quality control, exclusion of unwanted events and adjustment of compensation, biologically relevant gates are set. The gate definitions are then exported as a FlowJo Workspace file or GatingML file from FlowJo or Cytobank, respectively. The FlowJo or GatingML file is then imported into MetaGate, which parses the file and produces a list of defined gates (Figure 1B). In the MetaGate graphical user interface, the user can then define populations by combining the gates, e.g. defining “CD8 T cells” as events inside the “CD3+” and “CD8+” gate, but outside the “CD19+” gate. The MetaGate data reduction algorithm is then applied, using the definitions of gates and populations along with raw data from FCS files to calculate mean, median and geometric intensity values and frequencies of all populations in each population. Given *P* populations and *M* markers, the algorithm will output (3 * M + P) * P values for each sample. Assuming 100,000 events, 40 markers, 100 populations and 4 bytes per value, MetaGate will generate 86 KB of data from a 15 MB FCS file. This data is then stored as a data file that is used for all subsequent data analysis (Figure 1C).

**Figure 1.**
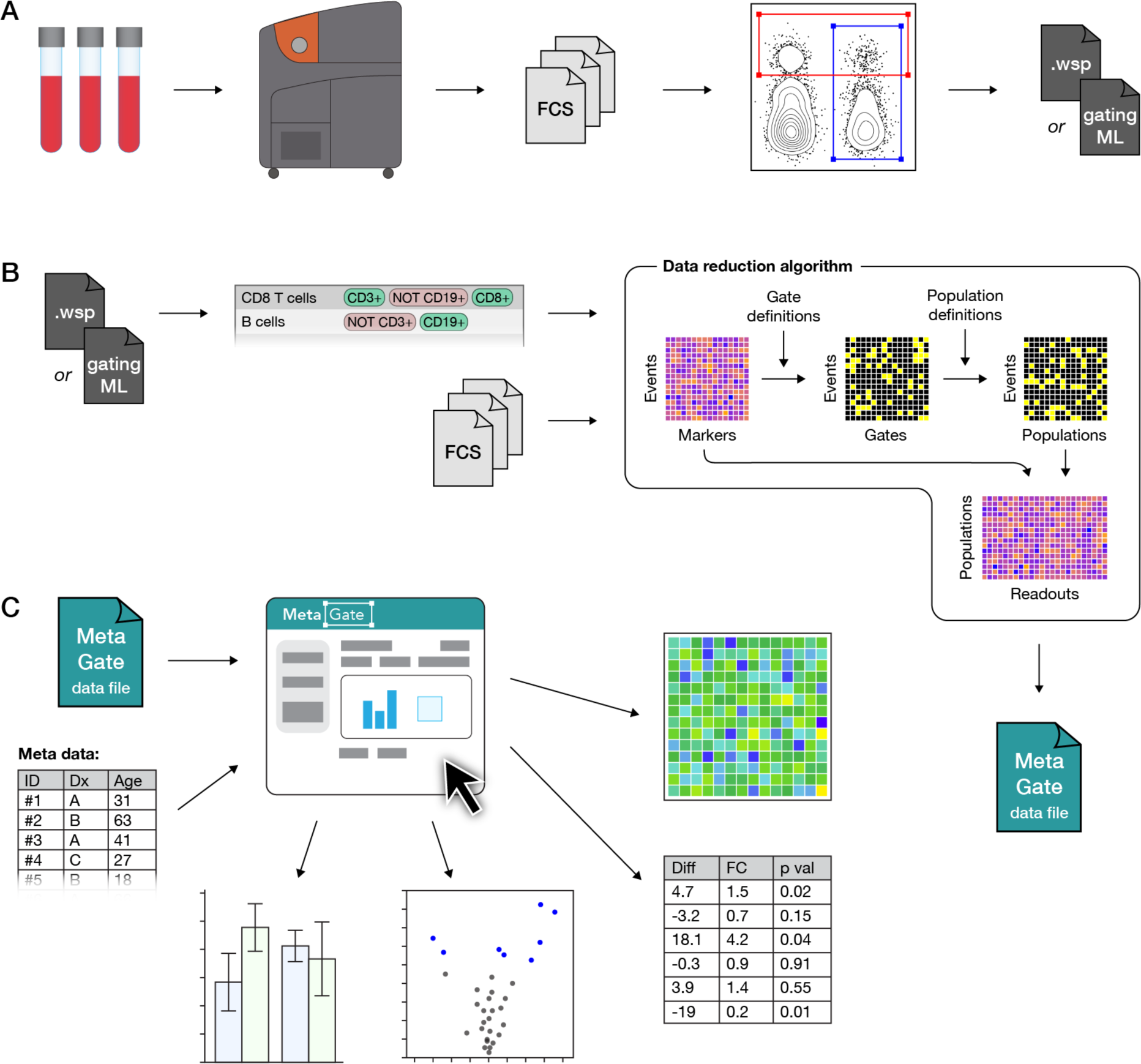
MetaGate data analysis workflow. (A) A biological sample, such as patient blood, is analyzed using a mass or flow cytometer, which produces FCS data files. Manual gating is performed in FlowJo or Cytobank, creating a data file with specifications of each gate. (B) Gate data and FCS files are imported into MetaGate, where a graphical user interface allows defining populations based on combinations of gates. Through a data reduction algorithm, a MetaGate data file is created, which contains marker expression and event frequencies of combinations of populations. (C) The self-containing MetaGate data file is opened in the MetaGate graphical user interface for interactive production of statistics and plots, such as heatmaps, volcano plots and bar plots.

### Data analysis in MetaGate

After loading the MetaGate data file in the MetaGate graphical user interface, the user can upload sample meta data, such as clinical features, experimental conditions or sample timepoints (Figure 1C). Sample *groups* are then defined interactively by selecting features based on the meta data.

The meta data should include information about which panel is used for each sample. By setting this as a *panel variable*, MetaGate will automatically make sure that the same individual is not included twice in a comparison in cases where both panels would provide the same data. In projects that contain paired samples, such as multiple perturbations or timepoints, a variable should be included that uniquely identifies each patient or healthy donor. MetaGate will then use this variable to perform paired statistical analyses. All meta data and group definitions are stored in the MetaGate file but can be modified at any time in downstream analysis.

To demonstrate the main features of MetaGate, a previously partially reported data set of immune cell characterization in diffuse large B-cell lymphoma (DLBCL) was analyzed. Peripheral blood mononuclear cells (PBMC) from a total of 28 DLBCL patients and 17 age- and sex-matched healthy controls (Table 1) were investigated using two mass cytometry panels (Figure 2, Supplementary Tables 1–2). To evaluate the effect of therapy, patients were sampled both at the time of diagnosis and after treatment with rituximab and chemotherapy. For each of the two panels separately, gating was performed in FlowJo. The two resulting MetaGate data files were then merged. All plots and statistical calculations in Figure 3–5 and accompanying supplementary tables were produced in MetaGate.

**Figure 2.**
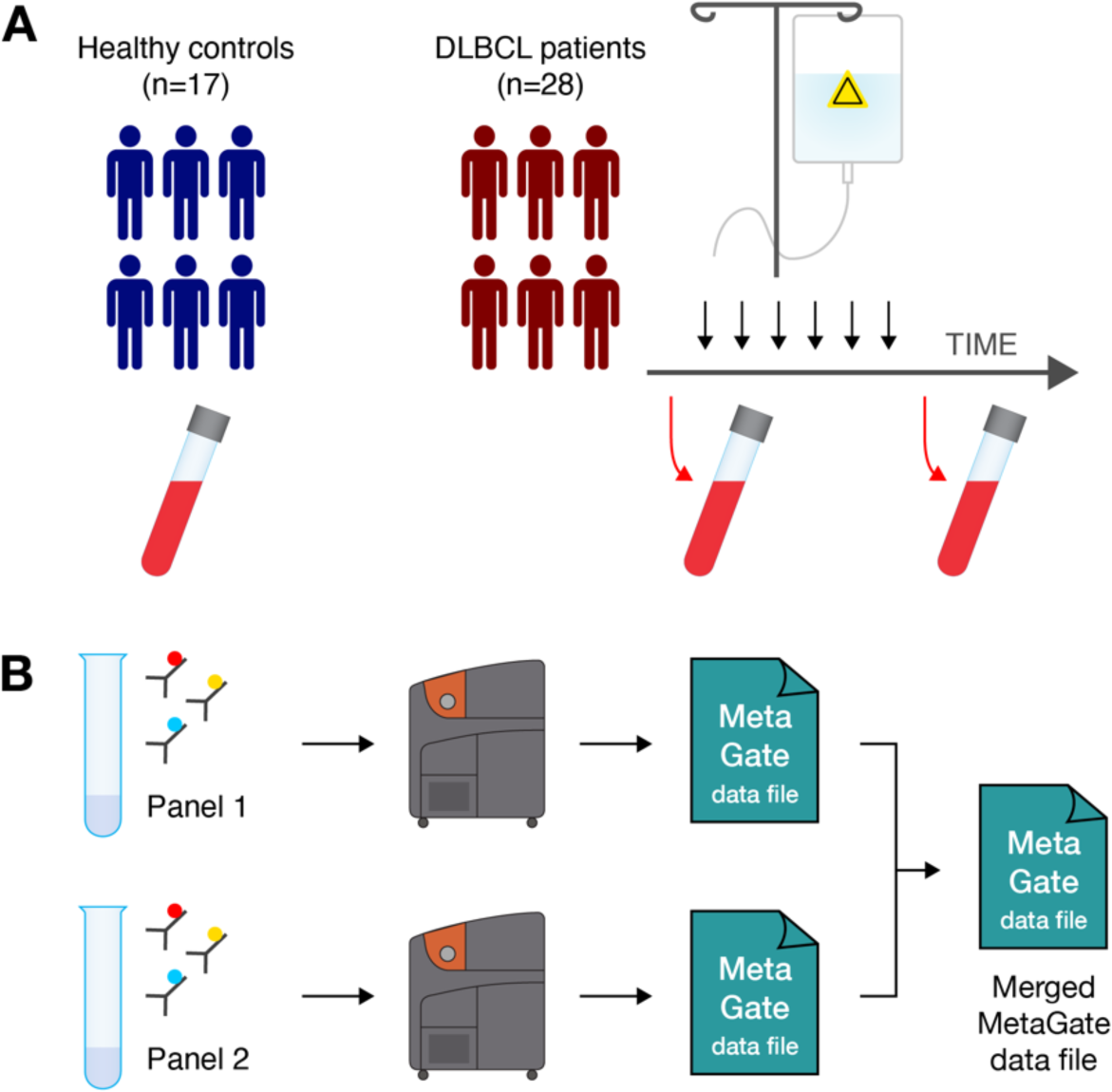
DLBCL immune characterization workflow. (A) Peripheral blood was collected from healthy blood donors (n=17) and from patients diagnosed with diffuse large B-cell lymphoma (n=28) before and after chemotherapy. (B) Blood samples were split and analyzed using two mass cytometry panels. Data from each panel was imported separately in MetaGate and later merged.

**Table 1.** Patients and healthy controls.

|  | Healthy controls | Patients |
| --- | --- | --- |
| Number of individuals | 17 | 28 |
| Female | 9 (53%) | 12 (43%) |
| Median age | 67 | 65 |
| *Subtype* |  |  |
| GCB DLBCL |  | 13 (46.4%) |
| Non-GCB DLBCL |  | 11 (39.3%) |
| Other |  | 4 (14.3%) |
| *Stage* |  |  |
| Stage I |  | 3 (10.7%) |
| Stage II |  | 7 (25%) |
| Stage III |  | 3 (10.7%) |
| Stage IV |  | 15 (53.6%) |

### Large impact of DLBCL on peripheral blood immune cell phenotypes

MetaGate allows creation of three main types of heatmaps. Using the first type, which shows marker expression for multiple populations in one group, the defining expression patterns of the key included populations can be visualized (Figure 3A).

**Figure 3.**
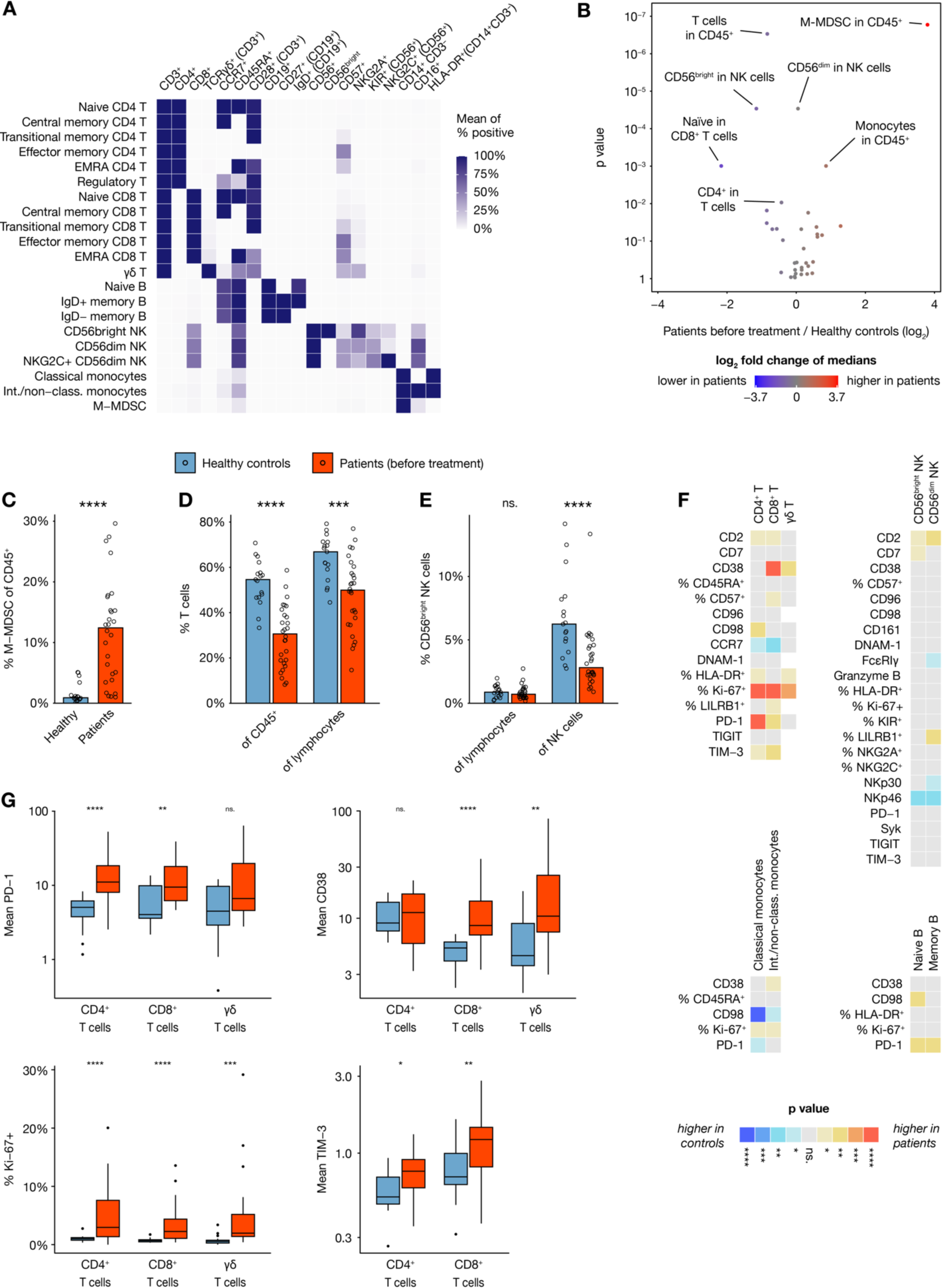
Peripheral blood immune cell composition in DLBCL. (A) Heat map showing expression of key markers in subsets of analyzed cell types, visualizing how subsets were defined for downstream analysis. (B) Volcano plot showing size differences of 36 key immune cell types between healthy donors and all patients before chemotherapy. (C– E) Bar plots showing percentages of (C) M–MDSC (defined as HLA-DR^-^ CD14^+^ CD19^-^ CD3^-^ CD56^-^ cells), (D) T cells and (E) CD56^bright^ NK cells, within various parent populations in healthy controls (n=17) and all patients before therapy (n=28). (F) Heatmap showing differences in marker expression between healthy controls (n=17) and patients before therapy (n=21–28) within multiple immune cell subsets, with colors indicating direction of difference and statistical significance from nonparametric tests without p value adjustment. Values are mean intensity values unless otherwise indicated. (G) Box plots showing selected readouts from (F).

Volcano plots are useful for quickly identifying main differences between two groups, as they provide a graphical representation of both statistical significance and magnitude of difference for multiple readouts in the same plot. In MetaGate, volcano plots can be generated based on data from multiple panels and explored interactively by holding the cursor over each dot. Using a volcano plot to compare sizes of major cell subsets between healthy donors and DLBCL patient samples before therapy, reveals multiple large differences (Figure 3B, Supplementary Table 5). Most significantly, HLA-DR^-^ CD14^+^ CD19^-^ CD3^-^ CD56^-^ cells, indicative of monocytic myeloid-derived suppressor cells,^21^ are greatly expanded in patients (Figure 3C). Inversely, the T-cell fraction of all CD45^+^ is lower in patients, but T cells also constitute a smaller fraction of lymphocytes (Figure 3D). As mass cytometry, in contrast to flow cytometry, does not allow distinction of lymphocytes by morphology, the lymphocyte population is here defined as the sum of T, B and natural killer (NK) cells. In patients, the CD56^bright^ cells constitute a smaller part of the NK cell compartment, relative to the more mature CD56^dim^ cells (Figure 3E).

The second main type of heatmaps that MetaGate can produce, enables two-group comparisons of multiple markers in multiple populations (Figure 3F). Markers can represent both marker intensities and percentages of positive cells, and data from multiple panels can be displayed in the same plot. Using colors for displaying the p values from multiple non-parametric tests and the direction of change, these plots give a fast overview of potentially significant findings. MetaGate furthermore produces a complete table of all statistics and allows this to be exported as a Microsoft Excel file. Most strikingly, T cells of DLBCL patients display higher levels of CD38, Ki-67, PD-1 and TIM-3 (Figure 3G).

### Immune cell subset dynamics through the course of treatment

In addition to slightly varying chemotherapy regimens, the anti-CD20 antibody rituximab was given to all patients. As expected, peripheral blood B cells were virtually non-detectable in post-treatment samples, while B cell numbers before treatment did not differ significantly from those of healthy controls (Figure 4A). As illustrated here, MetaGate automatically selects appropriate statistical tests based on the number of groups compared.

**Figure 4.**
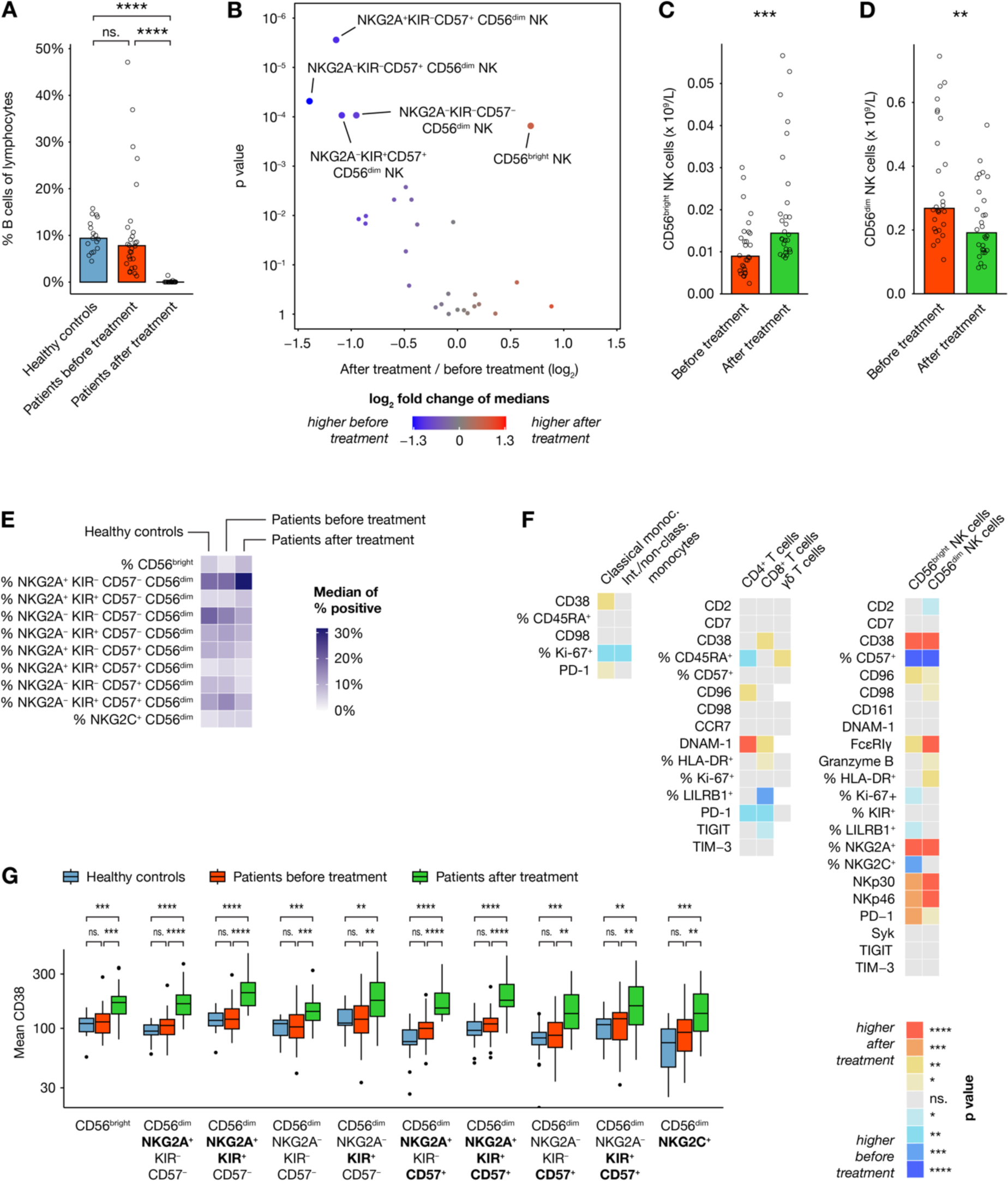
Effects of treatment on immune cell phenotypes. (A) B cell frequencies as percentage of all CD45^+^ in healthy controls (n=17) and all patients (n=28) before and after treatment. (B) Volcano plot showing differences in absolute counts of 28 cell subsets before and after treatment (n=28). (C–D) Selected comparisons from (B). (E) Heatmap showing median frequencies of key NK cell subsets as percentage of bulk NK cells in healthy controls (n=17) and patients (n=28) before and after therapy. (F) Heatmap showing differences in marker expression within multiple immune cell subsets between patients before and after treatment (n=20–28), with colors indicating direction of difference and statistical significance from paired nonparametric tests without p value adjustment. (G) Mean CD38 expression in multiple NK cell subsets of healthy controls (n=15–17) and patients (n=25–28) before and after treatment.

The observed B-cell depletion highlights the importance of assessing absolute cell counts, in contrast to the relative subset sizes usually provided by cytometry assays. If absolute counts of a population are available, MetaGate automatically calculates absolute counts of all subpopulations. By linking clinical lymphocyte counts to the lymphocyte population in MetaGate, absolute counts of key T, B and NK cell subsets could be assessed. Most significantly, patients displayed larger numbers of the CD56^bright^ NK cells after therapy, while several subsets of the more mature CD56^dim^ NK cells decreased in size (Figures 4B–D). The NK-cell subset dynamics can be further investigated by utilizing the third type of heatmap available in MetaGate, which allows visualization of multiple readouts across more than two groups (Figure 4E). In addition to the expansion of the CD56^bright^ NK cells, the CD56^dim^ compartment displays a shift towards less mature cells with more NKG2A-expressing and less CD57-expressing cells. Looking at changes in marker expression after therapy, this is corroborated by the observed increase in NKp30 and NKp46 expression (Figure 4F). Furthermore, a clear increase in CD38 expression is observed in NK cells, consistent across all major subsets (Figure 4G).

### Prediction of disease outcome

Using provided meta data, MetaGate allows simple and dynamic creation of sample groups for visualization and statistical testing. Looking at absolute cell counts of key lymphocyte populations in patient samples taken at the time of diagnosis, no clear differences were seen based on major age and subtype groups (Figure 5A–B). However, advanced disease (Ann Arbor stage III or IV) was somewhat associated with lower numbers of CD4^+^ T cells and CD56^bright^ NK cells (Figures 5C–E). Only five patients experienced disease progression during the follow-up time. Still, this group showed an association with lower absolute counts of CD56^dim^ NK cells and higher numbers of IgD^−^ memory B cells (Figures 5F–H).

**Figure 5.**
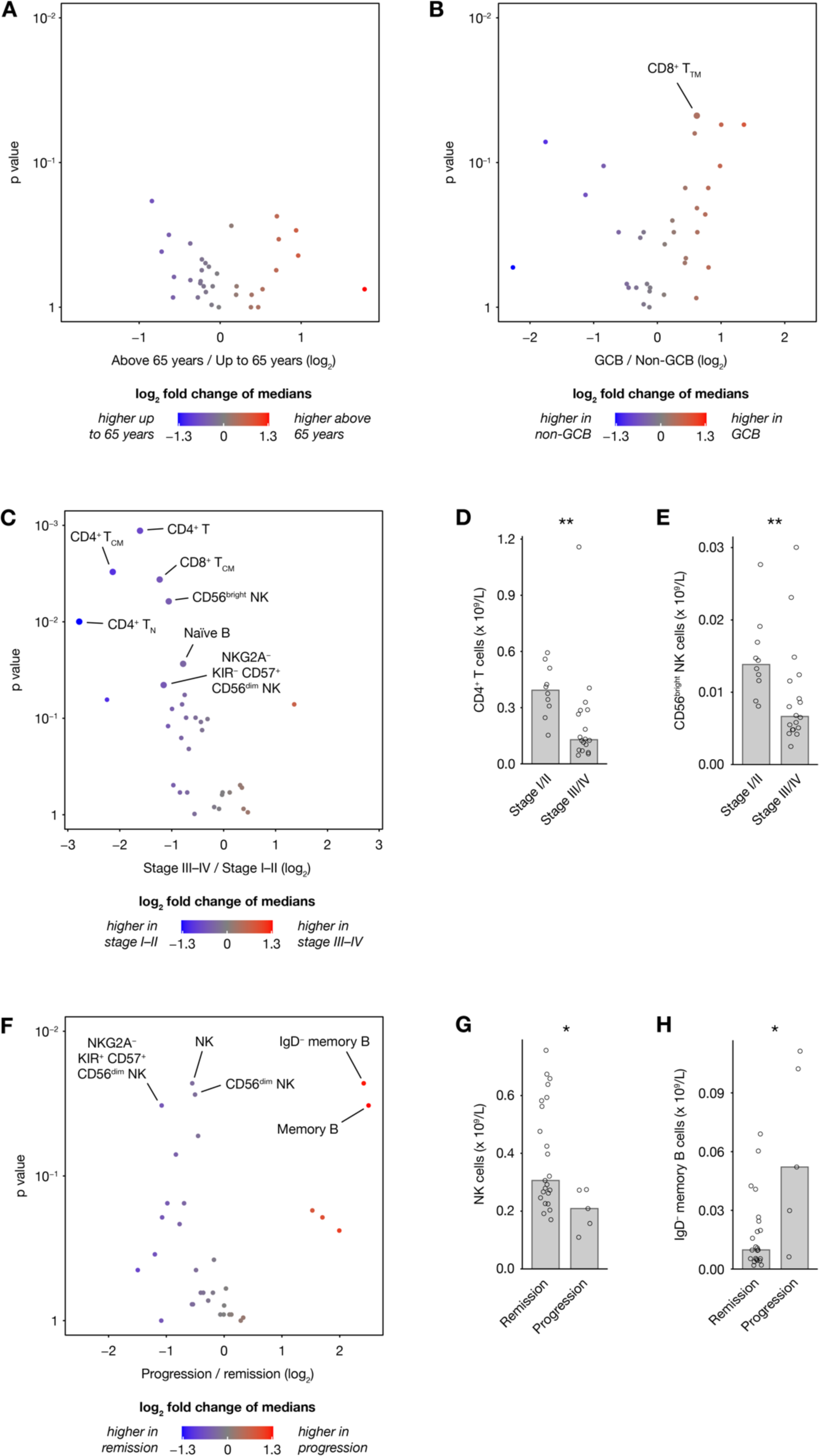
Immune cell repertoires stratified on patient characteristics. (A–C, F) Volcano plots showing differences in 33 absolute cell counts in peripheral blood of patients before therapy, stratified on (A) age, (B) subtype, (C) stage and (F) disease progression within the follow-up time. (D–E, G–H) Selected readouts from (C) and (F).

## Discussion

The continuously increasing complexity of cytometry data warrants new strategies for data analysis. We developed MetaGate, allowing interactive and fast statistical analysis and visualization of complex cytometry data sets. In this paper, we visualize the novel features of MetaGate through the analysis of a previously partly published broad multi-panel mass cytometric characterization of peripheral blood immune cells in a cohort of 28 DLBCL patients.

All plots and statistical analyses throughout this paper were generated in MetaGate, illustrating many of the most important features of the software package. Modern cytometry data sets often contain large numbers of readouts for comparison and assessing all of them manually can be very laborious, especially when there is a need to stratify the data on multiple clinical variables. Volcano plots, which are routinely used in genomics and proteomics, allow both statistical significance and the magnitude of change to be displayed in one graphical representation, which in MetaGate can be explored interactively. Conversely, heatmaps allow more than two groups to be compared, or multiple readouts to be assessed in multiple populations. Importantly, when comparing two groups, MetaGate heatmaps can also display statistical significance and direction of change, which can be particularly useful when assessing marker expression across multiple cell subsets. Such large-scale statistical testing introduces a considerable risk of type I errors. While MetaGate offers several p value correction techniques that can partly alleviate this problem, the use of p values in heatmaps and volcano plots in MetaGate should primarily be considered as a data exploration method, useful for highlighting potential findings of interest. Such findings can then be further explored using bar plots, which also allow multi-group comparisons and visualization of other meta data. In all plots, MetaGate automatically selects appropriate non-parametric statistical tests.

In cytometry experiments with clear groups of samples, for example perturbation and controls, resulting data from manual gating can relatively easily be managed manually for statistics and visualization. However, studies involving clinical data often include multiple variables of meta data, such as age, sex, diagnosis, sampling timepoint and treatment response. In this case, appropriate sample groups and comparisons may be numerous and not necessarily obvious early in the data analysis workflow. This can make manual data handling laborious and prone to errors. MetaGate seeks to alleviate this by mapping meta data from separate data files to samples and allowing groups to be created through a point-and-click query system in which the user selects features from the imported meta data. As both meta data and group definitions can be modified at any time, data exploration becomes simple and efficient.

All data analysis in MetaGate is based on manual gating of the data, meaning that cell types are defined by manually setting presumed biological relevant cut-offs for marker expression in several one- or two-dimensional data plots. Although remaining the most common data analysis strategy, manual gating has multiple drawbacks.^22^ The reliance on visual inspection of data by a trained professional introduces potential operator bias and confirmation bias. Furthermore, with the increasing complexity of cytometry data, manual gating represents a laborious analysis strategy. Many of the analysis algorithms developed in response to these challenges prove particularly useful for exploring novel or complex cell subsets, but may not produce results that are easily compared between different studies or experimental batches.^9^ *DeepCyTOF* and *flowLearn* are examples of algorithms that address these obstacles by automating the manual gating procedure through machine learning.^23, 24^ While MetaGate relies on gating of cells, there is no intrinsic requirement for these gates to be created manually by humans. Therefore, MetaGate can be further developed to allow (semi-)automatic gating by any of these algorithms upstream of the interactive statistical analysis in MetaGate.

The MetaGate data reduction algorithm works by calculating mean intensity values and sizes of all defined populations for each sample, producing a very condensed data set that can be used for downstream analysis without access to the raw data. Consequently, MetaGate can only generate plots and statistics based on predefined populations, limiting its usefulness for exploration of novel cell subsets. However, there are multiple benefits to this strategy. Because cytometry data consists of single-cell measurements of multiple parameters, data sets are typically large. A theoretical set of 100 files with one million events and 40 parameters in each would create around 15 gigabytes of data, which exceeds the available memory of most common workstations. Furthermore, the computational expensiveness of gating is increasing with the number of events and parameters. By performing all the memory- and processor-consuming tasks in the MetaGate data import procedure, the downstream analysis in MetaGate becomes comparably very fast. Fixing gates, population definitions and sample selections at one point, and making these visible to the user, also enhances the traceability of the analysis. This, and the small size of the data file, furthermore simplifies data sharing, making data analysis possible without in-depth experimental knowledge, powerful computers or access to other specialized software.

MetaGate is fully written in the R programming language, utilizing the *shiny*^15^ package to provide a web browser-based user interface. Taking advantage of the large selection of available R packages, the functionality of MetaGate can easily be extended. As a shiny-based application, MetaGate can either run locally on the user’s computer or be run on a remote server and accessed through the internet. As internet connection is not required and all source code is open and without need of compilation, MetaGate can also be used in secure data environments where custom software installation is prohibited, as long as R is available.

While demonstrating some of the most important features of MetaGate, the mass cytometry analysis of 28 DLBCL patients and matched controls reveals marked effects on the peripheral blood immune system of DLBCL patients. Although current therapy induces remission in a large majority of DLBCL patients, incomplete remission or relapses are seen in around one-third of the patients, and a better understanding of the immune responses could potentially lead to improved prognostication and treatment customization.^12^ Monocytic myeloid-dervied suppressor cells (M-MDSCs) are pathologically activated monocytes that have been associated with immunosuppression and poor outcome in multiple cancer settings.^25^ Our data shows high numbers of M-MDSCs among DLBCL patients, which has previously been reported and linked to immunosuppression,^26, 27^ potentially explaining why monocytosis was identified as a negative prognostic marker in DLBCL^28^. Furthermore, the increased expression of Ki-67, CD38, PD-1 and TIM-3 on T cells represents a phenotype consistent with exhaustion and potential dysfunctional activation.^29, 30^

Apart from the expected near-total depletion of B cells, the most markedly effect of chemotherapy on peripheral blood immune cell phenotypes was seen for NK cells. After chemotherapy, NK cells displayed lower expression of the maturation marker CD57, while higher expression was seen for the inhibitory receptor NKG2A and activating receptors NKp30 and NKp46, which is in line with observations of reconstitution of NK cell subsets after hematological stem cell transplantation.^31^ The broad upregulation of CD38 expression across all NK cell subsets suggests a systemic immune activation following chemo-immunotherapy, possibly reflecting homeostatic recovery. Corroborating previous DLBCL studies, our data showed a positive correlation between NK cell counts before initiation of therapy and beneficial outcome.^32, 33^

In conclusion, we present a new bioinformatical tool for high-throughput statistical analysis and visualization of cytometry data. The features of this software are displayed through the analysis of a mass cytometry characterization of peripheral blood from 28 DLBCL patients and matched controls, highlighting large immunophenotypic effects of both the disease and chemoimmunotherapy treatment, corroborating previously published reports. The initial manual gating of data, data reduction algorithm and dynamic integration with meta data, simplifies feature selection, data sharing and generation of publication-ready statistics and plots. Published as an open-source R package, MetaGate can be improved, customized and integrated in existing workflows, potentially allowing researchers to more easily tackle the continuously increasing complexity of cytometry data.

## Acknowledgements

The project was supported by The Research Council of Norway (Project number 275469, 237579), Center of Excellence: Precision Immunotherapy Alliance, 332727; The Norwegian Cancer Society (Project numbers 190386, 223310), The South-Eastern Norway Regional Health Authority (2021-073), EU H2020-MSCA Research and Innovation programme (Project number 801133), Knut and Alice Wallenberg Foundation, Swedish Foundation for Strategic Research, the US National Cancer Institute P01 CA111412, Subaward No: P009500901. This work was further supported by grants from the Swedish Research Council (223310), the Swedish Children’s Cancer Society (PR2020-1059), the Swedish Cancer Society (21-1793Pj), Sweden’s Innovation Agency, the Karolinska Institutet, and the Swiss Cancer League (A.T-P.; grant no. BIL KFS-3745-08-2015). We are grateful for the support from The Flow Cytometry Core Facility at Oslo University Hospital, Radiumhospitalet.

## Author contributions

A.T-P., H.J.H. and E.H.A. conducted experiments. E.H.A. conceptualized software with input from A.T-P. and K-J.M. E.H.A. wrote the code. A.K. and H.H. provided clinical samples and clinical data. E.H.A. wrote the manuscript. All authors edited the manuscript.

## Supplementary Figures

**Supplementary Figure 1.**
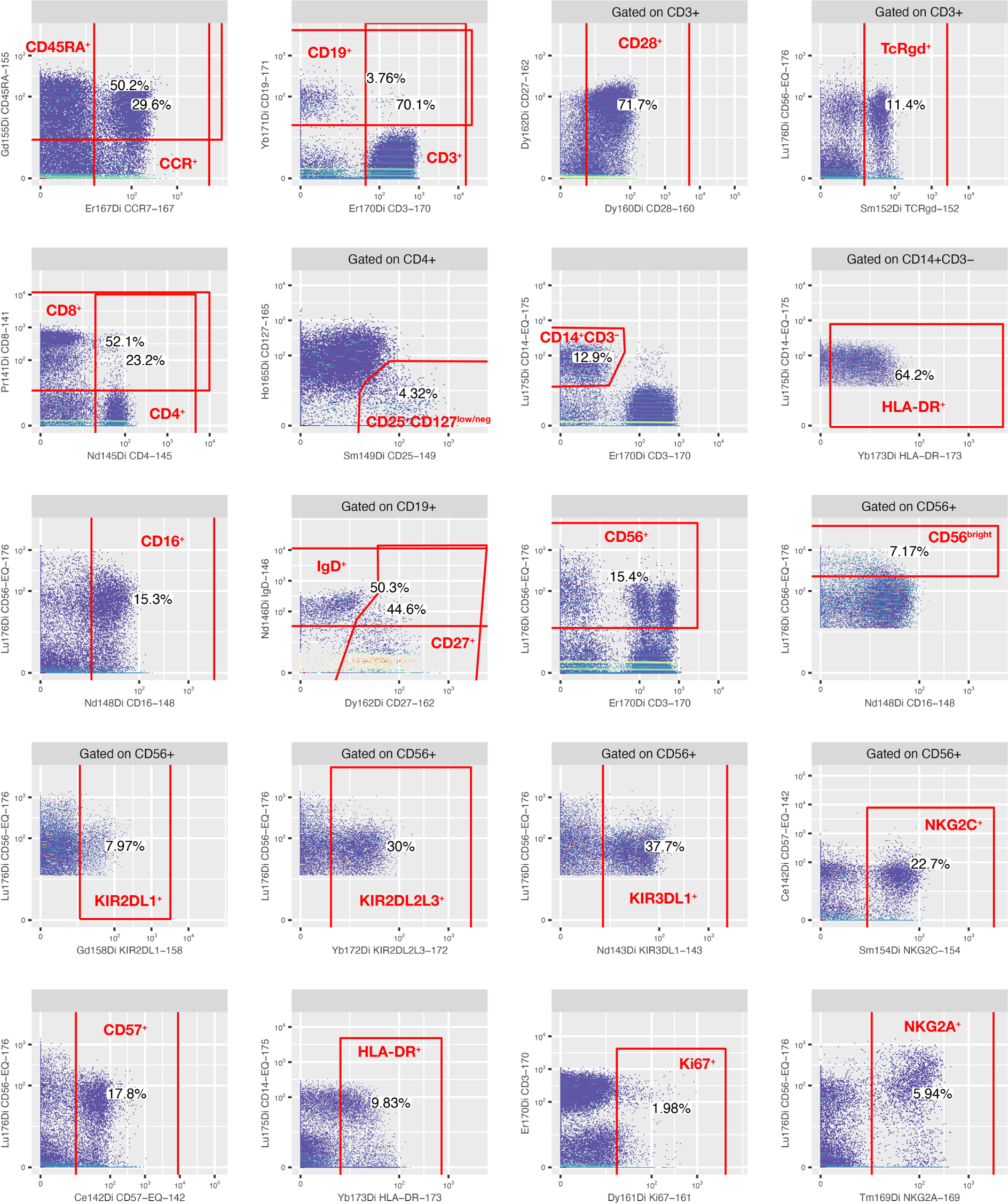
Gating strategy for panel 1.

**Supplementary Figure 2.**
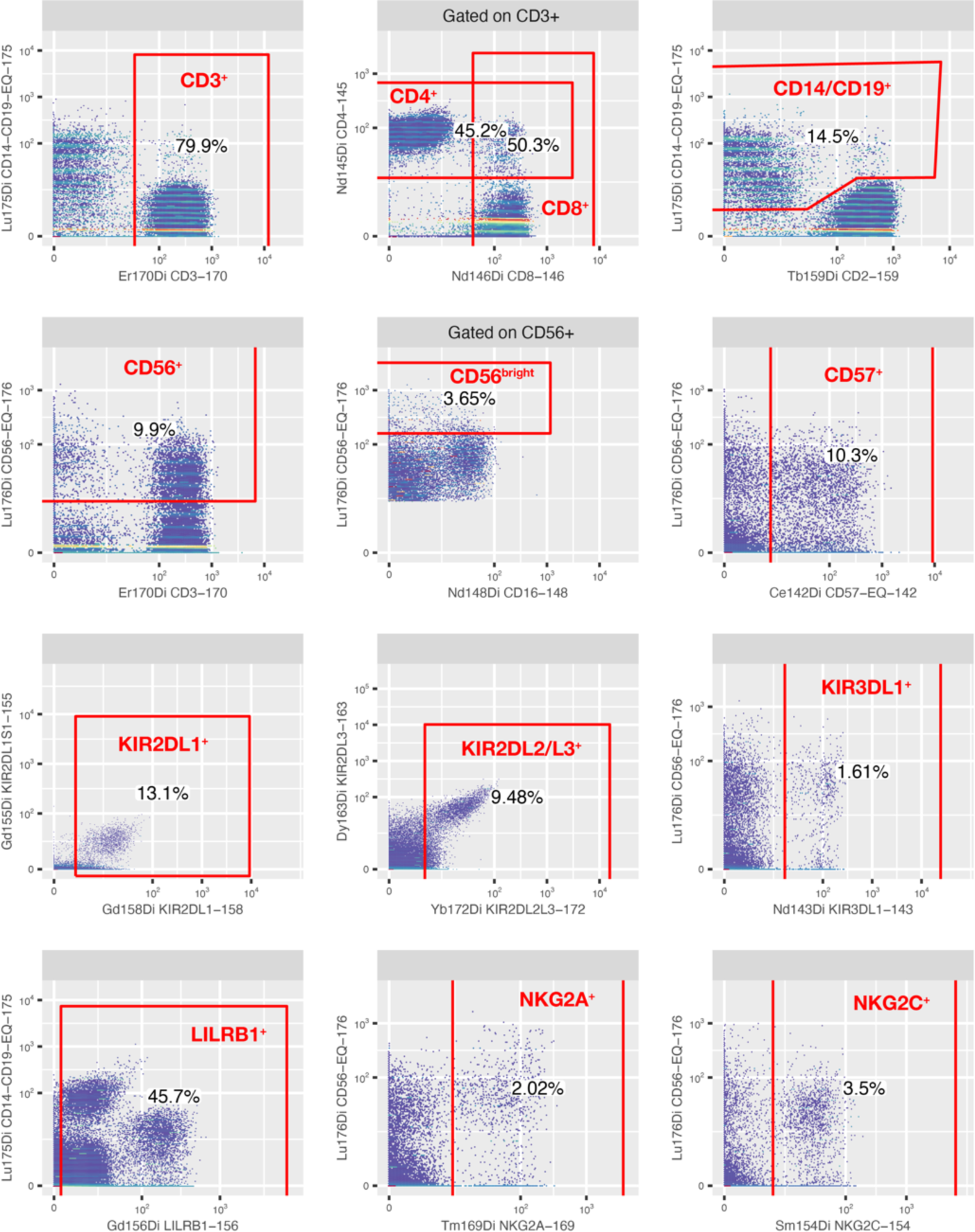
Gating strategy for panel 2.

## Supplementary Tables

**Supplementary Table 1.** Mass cytometry panel 1.

| Mass | Antigen | Clone | Supplier | Note |
| --- | --- | --- | --- | --- |
| 89Y | CD45 | HI30 | Fluidigm |  |
| 102Pd | Barcode |  | Fluidigm |  |
| 103Rh | Intercalator |  | Fluidigm |  |
| 104Pd | Barcode |  | Fluidigm |  |
| 105Pd | Barcode |  | Fluidigm |  |
| 106Pd | Barcode |  | Fluidigm |  |
| 108Pd | Barcode |  | Fluidigm |  |
| 110Pd | Barcode |  | Fluidigm |  |
| 141Pr | CD8 | RPA-T8 | Biolegend |  |
| 142Nd | CD57 | HCD57 | Fluidigm |  |
| 143Nd | KIR3DL1 | DX9 | Miltenyi |  |
| 144Nd | CD38 | REA572 | Miltenyi |  |
| 145Nd | CD4 | RPA-T4 | Fluidigm |  |
| 146Nd | IgD | IA6-2 | Fluidigm |  |
| 147Sm | CD71 | AC102 | Miltenyi | <sup>2</sup> |
| 148Nd | CD16 | 3G8 | Fluidigm |  |
| 149Sm | CD25 | 2A3 | Fluidigm |  |
| 150Nd | Anti-GFP, GLUT-1-GFP | FM264G | Biolegend | <sup>2</sup> |
| 151Eu | CD123 | AC145 | Miltenyi | <sup>1</sup> |
| 152Sm | TCRgd | 11F2 | Fluidigm |  |
| 153Eu | CD7 | CD7-6B7 | Fluidigm |  |
| 154Sm | NKG2C | REA205 | Miltenyi |  |
| 155Gd | CD45RA | HI100 | Fluidigm |  |
| 156Gd | NKp46 | 9E2 | Miltenyi |  |
| 158Gd | KIR2DL1 | REA284 | Miltenyi |  |
| 159Tb | CD2 | RPA-2.10 | eBioscience |  |
| 160Gd | CD28 | CD28.2 | Fluidigm |  |
| 161Dy | Ki67 | B56 | Fluidigm |  |
| 162Dy | CD27 | L128 | Fluidigm |  |
| 163Dy | CD98 | REA387 | Miltenyi |  |
| 164Dy | CD161 | HP-3G10 | Fluidigm |  |
| 165Ho | CD127 | A019D5 | Fluidigm |  |
| 166Er | CD11c | B-ly6 | BD | <sup>1</sup> |
| 167Er | CCR7 | G043H7 | Fluidigm |  |
| 168Er | NKp30 | P30-15 | Miltenyi |  |
| 169Tm | NKG2A | Z199 | Fluidigm |  |
| 170Er | CD3 | UCHT1 | Fluidigm |  |
| 171Yb | CD19 | Æ1 | In-house |  |
| 172Yb | KIR2DL2L3 | GL183 | Miltenyi |  |
| 173Yb | HLA-DR | AC122 | Miltenyi |  |
| 174Yb | PD-1 | EH12.2H7 | Fluidigm |  |
| 175Lu | CD14 | M5E2 | Fluidigm |  |
| 176Yb | CD56 | NCAM16.2 | Fluidigm |  |
| 191/193Ir | Intercalator |  | Fluidigm |  |

<sup>1</sup> Excluded because the marker is not of relevance to this analysis.
<sup>2</sup> Excluded due to batch effects.

**Supplementary Table 2.**
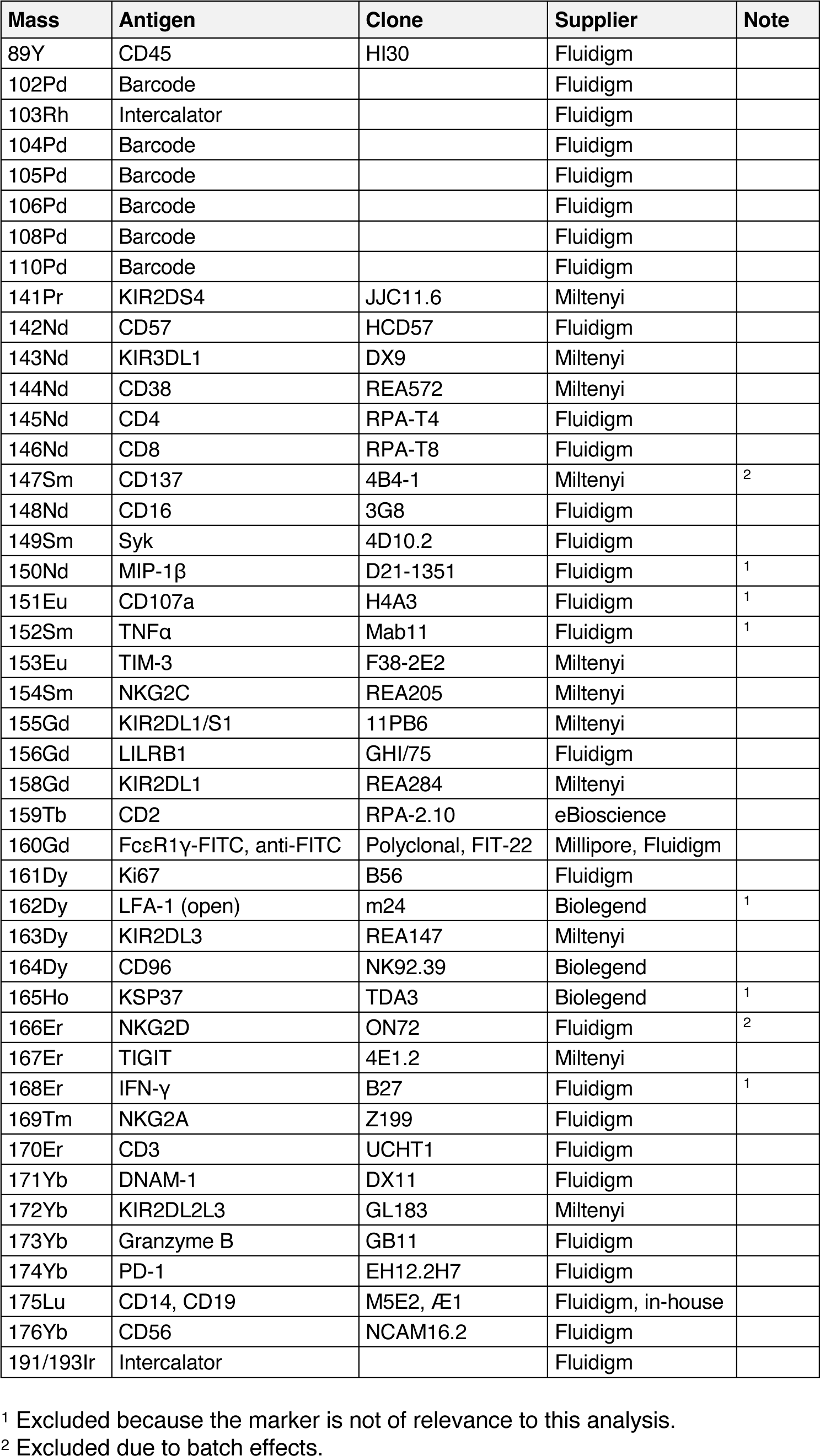
Mass cytometry panel 2.

**Supplementary Table 3.** Population definitions for panel 1.

| Name | Definition |
| --- | --- |
| Lymphocytes | Lymphocytes |
| T cells | CD3+, NOT CD19+, NOT CD14+CD3-, NOT CD56+ |
| CD4+ T cells | CD4+, NOT CD8+, CD3+, NOT CD19+, NOT CD14+CD3-, NOT CD56+ |
| Naive CD4+ T cells | CCR7+, CD45RA+, CD4+, NOT CD8+, CD3+, NOT CD19+, NOT CD14+CD3-, NOT CD56+ |
| Central memory CD4+ T cells | CCR7+, NOT CD45RA+, CD4+, NOT CD8+, CD3+, NOT CD19+, NOT CD14+CD3-, NOT CD56+ |
| Transitional memory CD4+ T cells | CD3+ => CD28+, NOT CCR7+, NOT CD45RA+, CD4+, NOT CD8+, CD3+, NOT CD19+, NOT CD14+CD3-, NOT CD56+ |
| Effector memory CD4+ T cells | NOT CD3+ => CD28+, NOT CCR7+, NOT CD45RA+, CD4+, NOT CD8+, CD3+, NOT CD19+, NOT CD14+CD3-, NOT CD56+ |
| EMRA CD4+ T cells | NOT CCR7+, CD45RA+, CD4+, NOT CD8+, CD3+, NOT CD19+, NOT CD14+CD3-, NOT CD56+ |
| Regulatory T cells | CD4+ => CD25+CD127lowneg, NOT CD8+, CD3+, NOT CD19+, NOT CD14+CD3-, NOT CD56+ |
| CD8+ T cells | CD8+, NOT CD4+, CD3+, NOT CD19+, NOT CD14+CD3-, NOT CD56+ |
| Naive CD8+ T cells | CCR7+, CD45RA+, CD8+, NOT CD4+, CD3+, NOT CD19+, NOT CD14+CD3-, NOT CD56+ |
| Central memory CD8+ T cells | CCR7+, NOT CD45RA+, CD8+, NOT CD4+, CD3+, NOT CD19+, NOT CD14+CD3-, NOT CD56+ |
| Transitional memory CD8+ T cells | CD3+ => CD28+, NOT CCR7+, NOT CD45RA+, CD8+, NOT CD4+, CD3+, NOT CD19+, NOT CD14+CD3-, NOT CD56+ |
| Effector memory CD8+ T cells | NOT CD3+ => CD28+, NOT CCR7+, NOT CD45RA+, CD8+, NOT CD4+, CD3+, NOT CD19+, NOT CD14+CD3-, NOT CD56+ |
| EMRA CD8+ T cells | NOT CCR7+, CD45RA+, CD8+, NOT CD4+, CD3+, NOT CD19+, NOT CD14+CD3-, NOT CD56+ |
| gd T cells | CD3+ => TCRgd+, NOT CD8+, NOT CD4+, CD3+, NOT CD19+, NOT CD14+CD3-, NOT CD56+ |
| B cells | CD19+, NOT CD3+, NOT CD56+, NOT CD14+CD3- |
| Naive B cells | NOT CD19+ => CD27+, CD19+, NOT CD3+, NOT CD56+, NOT CD14+CD3- |
| Memory B cells | CD19+ => CD27+, NOT CD3+, NOT CD56+, NOT CD14+CD3- |
| IgD+ memory B cells | CD19+ => IgD+, CD19+ => CD27+, NOT CD3+, NOT CD56+, NOT CD14+CD3- |
| IgD- memory B cells | NOT CD19+ => IgD+, CD19+ => CD27+, NOT CD3+, NOT CD56+, NOT CD14+CD3- |
| NK cells | CD56+, NOT CD3+, NOT CD14+CD3-, NOT CD19+ |
| CD56dim NK cells | NOT CD56+ => CD56bright, CD56+, NOT CD3+, NOT CD14+CD3-, NOT CD19+ |
| NKG2A+ KIR+ CD57+ CD56dim NK cells | NKG2A+, CD56+ => KIR+, CD57+, NOT CD56+ => CD56bright, CD56+, NOT CD3+, NOT CD14+CD3-, NOT CD19+ |
| NKG2A- KIR+ CD57+ CD56dim NK cells | NOT NKG2A+, CD56+ => KIR+, CD57+, NOT CD56+ => CD56bright, CD56+, NOT CD3+, NOT CD14+CD3-, NOT CD19+ |
| NKG2A+ KIR- CD57+ CD56dim NK cells | NKG2A+, NOT CD56+ => KIR+, CD57+, NOT CD56+ => CD56bright, CD56+, NOT CD3+, NOT CD14+CD3-, NOT CD19+ |
| NKG2A- KIR- CD57+ CD56dim NK cells | NOT NKG2A+, NOT CD56+ => KIR+, CD57+, NOT CD56+ => CD56bright, CD56+, NOT CD3+, NOT CD14+CD3-, NOT CD19+ |
| NKG2A+ KIR+ CD57- CD56dim NK cells | NKG2A+, CD56+ => KIR+, NOT CD57+, NOT CD56+ => CD56bright, CD56+, NOT CD3+, NOT CD14+CD3-, NOT CD19+ |
| NKG2A- KIR+ CD57- CD56dim NK cells | NOT NKG2A+, CD56+ => KIR+, NOT CD57+, NOT CD56+ => CD56bright, CD56+, NOT CD3+, NOT CD14+CD3-, NOT CD19+ |
| NKG2A+ KIR- CD57- CD56dim NK cells | NKG2A+, NOT CD56+ => KIR+, NOT CD57+, NOT CD56+ => CD56bright, CD56+, NOT CD3+, NOT CD14+CD3-, NOT CD19+ |
| NKG2A- KIR- CD57- CD56dim NK cells | NOT NKG2A+, NOT CD56+ => KIR+, NOT CD57+, NOT CD56+ => CD56bright, CD56+, NOT CD3+, NOT CD14+CD3-, NOT CD19+ |
| NKG2C+ CD56dim NK cells | CD56+ => NKG2C+, NOT CD56+ => CD56bright, CD56+, NOT CD3+, NOT CD14+CD3-, NOT CD19+ |
| CD56bright NK cells | CD56+ => CD56bright, NOT CD3+, NOT CD14+CD3-, NOT CD19+ |
| Monocytes | CD14+CD3- => HLA-DR+, NOT CD19+, NOT CD56+ |
| Classical monocytes | NOT CD16+, CD14+CD3- => HLA-DR+, NOT CD19+, NOT CD56+ |
| Int./non-class. monocytes | CD16+, CD14+CD3- => HLA-DR+, NOT CD19+, NOT CD56+ |
| M-MDSC | CD14+CD3-, NOT CD14+CD3- => HLA-DR+, NOT CD19+, NOT CD56+ |
| CCR7+ | CCR7+ |
| CD3+ | CD3+ |
| CD28+ (CD3+) | CD3+ => CD28+ |
| TCRgd+ (CD3+) | CD3+ => TCRgd+ |
| CD4+ | CD4+ |
| CD25+ CD127low/neg (CD4+) | CD4+ => CD25+CD127lowneg |
| CD8+ | CD8+ |
| CD14+ CD3- | CD14+CD3- |
| HLA-DR+ (CD14+ CD3-) | CD14+CD3- => HLA-DR+ |
| CD16+ | CD16+ |
| CD19+ | CD19+ |
| CD27+ (CD19+) | CD19+ => CD27+ |
| IgD+ (CD19+) | CD19+ => IgD+ |
| CD45RA+ | CD45RA+ |
| CD56+ | CD56+ |
| CD56bright | CD56+ => CD56bright |
| KIR+ (CD56+) | CD56+ => KIR+ |
| KIR2DL1+ CD56+ | CD56+ => KIR2DL1+ |
| KIR2DL2L3+ (CD56+) | CD56+ => KIR2DL2L3+ |
| KIR3DL1+ (CD56+) | CD56+ => KIR3DL1+ |
| NKG2C+ (CD56+) | CD56+ => NKG2C+ |
| CD57+ | CD57+ |
| HLA-DR+ | HLA-DR+ |
| Ki-67+ | Ki-67+ |
| NKG2A+ | NKG2A+ |

**Supplementary Table 4.** Population definitions for panel 2.

| Name | Definition |
| --- | --- |
| T cells | CD3+, NOT CD14+ or CD19+, NOT CD56+ |
| CD4+ T cells | CD3+ => CD4+, NOT CD3+ => CD8+, CD3+, NOT CD14+ or CD19+, NOT CD56+ |
| CD8+ T cells | CD3+ => CD8+, NOT CD3+ => CD4+, CD3+, NOT CD14+ or CD19+, NOT CD56+ |
| NK cells | CD56+, NOT CD3+, NOT CD14+ or CD19+ |
| CD56dim NK cells | NOT CD56+ => CD56bright, CD56+, NOT CD3+, NOT CD14+ or CD19+ |
| NKG2A+ KIR+ CD57+ CD56dim NK cells | NKG2A+, KIR+, CD57+, NOT CD56+ => CD56bright, CD56+, NOT CD3+, NOT CD14+ or CD19+ |
| NKG2A- KIR+ CD57+ CD56dim NK cells | NOT NKG2A+, KIR+, CD57+, NOT CD56+ => CD56bright, CD56+, NOT CD3+, NOT CD14+ or CD19+ |
| NKG2A+ KIR- CD57+ CD56dim NK cells | NKG2A+, NOT KIR+, CD57+, NOT CD56+ => CD56bright, CD56+, NOT CD3+, NOT CD14+ or CD19+ |
| NKG2A- KIR- CD57+ CD56dim NK cells | NOT NKG2A+, NOT KIR+, CD57+, NOT CD56+ => CD56bright, CD56+, NOT CD3+, NOT CD14+ or CD19+ |
| NKG2A+ KIR+ CD57- CD56dim NK cells | NKG2A+, KIR+, NOT CD57+, NOT CD56+ => CD56bright, CD56+, NOT CD3+, NOT CD14+ or CD19+ |
| NKG2A- KIR+ CD57- CD56dim NK cells | NOT NKG2A+, KIR+, NOT CD57+, NOT CD56+ => CD56bright, CD56+, NOT CD3+, NOT CD14+ or CD19+ |
| NKG2A+ KIR- CD57- CD56dim NK cells | NKG2A+, NOT KIR+, NOT CD57+, NOT CD56+ => CD56bright, CD56+, NOT CD3+, NOT CD14+ or CD19+ |
| NKG2A- KIR- CD57- CD56dim NK cells | NOT NKG2A+, NOT KIR+, NOT CD57+, NOT CD56+ => CD56bright, CD56+, NOT CD3+, NOT CD14+ or CD19+ |
| NKG2C+ CD56dim NK cells | NKG2C+, NOT CD56+ => CD56bright, CD56+, NOT CD3+, NOT CD14+ or CD19+ |
| CD56bright NK cells | CD56+ => CD56bright, NOT CD3+, NOT CD14+ or CD19+ |
| CD3+ | CD3+ |
| CD4+ | CD3+ => CD4+ |
| CD8+ | CD3+ => CD8+ |
| CD14+ or CD19+ | CD14+ or CD19+ |
| CD56+ | CD56+ |
| CD56bright | CD56+ => CD56bright |
| CD57+ | CD57+ |
| KIR+ | KIR+ |
| KIR2DL1+ | KIR2DL1+ |
| KIR2DL2L3+ | KIR2DL2L3+ |
| KIR3DL1+ | KIR3DL1+ |
| LILRB1+ | LILRB1+ |
| NKG2A+ | NKG2A+ |
| NKG2C+ | NKG2C+ |

**Supplementary Table 5.** Peripheral blood immune cell abundances in healthy donors and patients before therapy.

| Readout | Population | Median value |  | Subject count |  | log2 FC | p value |
| --- | --- | --- | --- | --- | --- | --- | --- |
|  |  | Healthy | Patients | Healthy | Patients |  |  |
| % T cells | Bulk | 54.63 % | 30.61 % | 17 | 28 | -0,8358211 | 2,96E-07 |
| % B cells | Bulk | 8.58 % | 4.76 % | 17 | 28 | -0,8504041 | 0,01527325 |
| % NK cells | Bulk | 11.74 % | 13.63 % | 17 | 28 | 0,21558804 | 0,15823181 |
| % Monocytes | Bulk | 8.26 % | 15.01 % | 17 | 28 | 0,86163219 | 0,00098764 |
| % M-MDSC | Bulk | 0.9 % | 12.43 % | 17 | 28 | 3,79518518 | 1,68E-07 |
| % CD4+ T cells | T cells | 55.73 % | 41.48 % | 17 | 28 | -0,4261842 | 0,00931575 |
| % CD8+ T cells | T cells | 39.18 % | 49.8 % | 17 | 28 | 0,34586705 | 0,01748529 |
| % gd T cells | T cells | 1.77 % | 1.31 % | 17 | 28 | -0,4353878 | 0,68515824 |
| % Naive CD4+ T cells | CD4+ T cells | 33.25 % | 18.42 % | 17 | 28 | -0,8518818 | 0,03309576 |
| % Central memory CD4+ T cells | CD4+ T cells | 32.55 % | 22.05 % | 17 | 28 | -0,5619033 | 0,0471276 |
| % Transitional memory CD4+ T cells | CD4+ T cells | 25.39 % | 42.61 % | 17 | 28 | 0,74687297 | 0,06931965 |
| % Effector memory CD4+ T cells | CD4+ T cells | 2.08 % | 3.18 % | 17 | 28 | 0,61297435 | 0,07700709 |
| % EMRA CD4+ T cells | CD4+ T cells | 1.05 % | 2.57 % | 17 | 28 | 1,28319028 | 0,03959916 |
| % Regulatory T cells | CD4+ T cells | 4.47 % | 5.68 % | 17 | 28 | 0,34762981 | 0,4649239 |
| % Naive CD8+ T cells | CD8+ T cells | 18.92 % | 4.21 % | 17 | 28 | -2,1674409 | 0,00098764 |
| % Central memory CD8+ T cells | CD8+ T cells | 5.89 % | 5.95 % | 17 | 28 | 0,01480148 | 0,37131182 |
| % Transitional memory CD8+ T cells | CD8+ T cells | 20.31 % | 22.48 % | 17 | 28 | 0,14646699 | 0,77219008 |
| % Effector memory CD8+ T cells | CD8+ T cells | 7.5 % | 11.44 % | 17 | 28 | 0,60868413 | 0,06571372 |
| % EMRA CD8+ T cells | CD8+ T cells | 30.48 % | 45.76 % | 17 | 28 | 0,58592034 | 0,04198928 |
| % Naive B cells | B cells | 83.76 % | 81.87 % | 17 | 28 | -0,0328609 | 0,58595842 |
| % Memory B cells | B cells | 16.24 % | 18.13 % | 17 | 28 | 0,15853257 | 0,58595842 |
| % IgD+ memory B cells | Memory B cells | 21.84 % | 13.52 % | 17 | 23 | -0,6915886 | 0,04803052 |
| % IgD- memory B cells | Memory B cells | 78.16 % | 86.48 % | 17 | 23 | 0,14588778 | 0,04803052 |
| % CD56bright NK cells | NK cells | 6.24 % | 2.81 % | 17 | 28 | -1,1521417 | 2,91E-05 |
| % CD56dim NK cells | NK cells | 93.76 % | 97.19 % | 17 | 28 | 0,05189189 | 2,91E-05 |
| % NKG2A+ KIR+ CD57+ CD56dim NK cells | CD56dim NK cells | 3.49 % | 4.82 % | 17 | 28 | 0,46470857 | 0,71955557 |
| % NKG2A- KIR+ CD57+ CD56dim NK cells | CD56dim NK cells | 12.96 % | 16.15 % | 17 | 28 | 0,31661505 | 0,75451763 |
| % NKG2A+ KIR- CD57+ CD56dim NK cells | CD56dim NK cells | 10.44 % | 9.38 % | 17 | 28 | -0,1539872 | 0,91701338 |
| % NKG2A- KIR- CD57+ CD56dim NK cells | CD56dim NK cells | 8.62 % | 8.53 % | 17 | 28 | -0,0149587 | 0,73697003 |
| % NKG2A+ KIR+ CD57- CD56dim NK cells | CD56dim NK cells | 4.52 % | 4.37 % | 17 | 28 | -0,0461083 | 0,88036026 |
| % NKG2A- KIR+ CD57- CD56dim NK cells | CD56dim NK cells | 10.74 % | 11.97 % | 17 | 28 | 0,15730461 | 0,50865543 |
| % NKG2A+ KIR- CD57- CD56dim NK cells | CD56dim NK cells | 19.81 % | 19.32 % | 17 | 28 | -0,0358563 | 0,93540976 |
| % NKG2A- KIR- CD57- CD56dim NK cells | CD56dim NK cells | 22.08 % | 16.9 % | 17 | 28 | -0,385279 | 0,09440922 |
| % NKG2C+ CD56dim NK cells | CD56dim NK cells | 3.56 % | 5 % | 17 | 28 | 0,48672765 | 0,35894261 |
| % Classical monocytes | Monocytes | 87.76 % | 85.08 % | 17 | 28 | -0,0448749 | 0,38393621 |
| % Int./non-class. monocytes | Monocytes | 12.24 % | 14.92 % | 17 | 28 | 0,28647881 | 0,38393621 |

## References

1. Bendall SC, Nolan GP, Roederer M, Chattopadhyay PK. A deep profiler’s guide to cytometry. Trends Immunol. 2012;33(7):323–32.

2. Grant R, Coopman K, Medcalf N, Silva-Gomes S, Campbell JJ, Kara B, et al. Quantifying Operator Subjectivity within Flow Cytometry Data Analysis as a Source of Measurement Uncertainty and the Impact of Experience on Results. PDA J Pharm Sci Technol. 2021;75(1):33–47.

3. Lugli E, Roederer M, Cossarizza A. Data analysis in flow cytometry: the future just started. Cytometry A. 2010;77(7):705–13.

4. Cheung M, Campbell JJ, Whitby L, Thomas RJ, Braybrook J, Petzing J. Current trends in flow cytometry automated data analysis software. Cytometry A. 2021;99(10):1007–21.

5. Van der Maaten L, Hinton G. Visualizing data using t-SNE. Journal of machine learning research. 2008;9(11).

6. Levine JH, Simonds EF, Bendall SC, Davis KL, Amir el AD, Tadmor MD, et al. Data-Driven Phenotypic Dissection of AML Reveals Progenitor-like Cells that Correlate with Prognosis. Cell. 2015;162(1):184–97.

7. Qiu P, Simonds EF, Bendall SC, Gibbs KD, Jr., Bruggner RV, Linderman MD, et al. Extracting a cellular hierarchy from high-dimensional cytometry data with SPADE. Nat Biotechnol. 2011;29(10):886–91.

8. Van Gassen S, Callebaut B, Van Helden MJ, Lambrecht BN, Demeester P, Dhaene T, et al. FlowSOM: Using self-organizing maps for visualization and interpretation of cytometry data. Cytometry A. 2015;87(7):636–45.

9. Newell EW, Cheng Y. Mass cytometry: blessed with the curse of dimensionality. Nat Immunol. 2016;17(8):890–5.

10. Morton LM, Wang SS, Devesa SS, Hartge P, Weisenburger DD, Linet MS. Lymphoma incidence patterns by WHO subtype in the United States, 1992-2001. Blood. 2006;107(1):265-76.

11. Alizadeh AA, Eisen MB, Davis RE, Ma C, Lossos IS, Rosenwald A, et al. Distinct types of diffuse large B-cell lymphoma identified by gene expression profiling. Nature. 2000;403(6769):503-11.

12. Sehn LH, Salles G. Diffuse Large B-Cell Lymphoma. N Engl J Med. 2021;384(9):842–58.

13. Ask EH, Tschan-Plessl A, Gjerdingen TJ, Saetersmoen ML, Hoel HJ, Wiiger MT, et al. A Systemic Protein Deviation Score Linked to PD-1(+) CD8(+) T Cell Expansion That Predicts Overall Survival in Diffuse Large B Cell Lymphoma. Med (N Y). 2021;2(2):180–95 e5.

14. R Core Team. R: A language and environment for statistical computing.: R Foundation for Statistical Computing, Vienna, Austria.; 2020.

15. Chang. W, Cheng. J, Allaire. J, Sievert. C, Schloerke. B, Xie. Y, et al. shiny: Web Application Framework for R. R package version 1.6.0 ed2021.

16. Finak G, Jiang M. flowWorkspace: Infrastructure for representing and interacting with gated and ungated cytometry data sets. R package version 4.2.0 ed2020.

17. Finak G, Jiang W, Gottardo R. CytoML for cross-platform cytometry data sharing. Cytometry A. 2018;93(12):1189–96.

18. Hahne F, LeMeur N, Brinkman RR, Ellis B, Haaland P, Sarkar D, et al. flowCore: a Bioconductor package for high throughput flow cytometry. BMC Bioinformatics. 2009;10:106.

19. Spidlen J, Gopalakrishnan N, Hahne F, Ellis B, Gentleman R, Dalphin M, et al. flowUtils: Utilities for flow cytometry. R package version 1.54.0 ed2020.

20. Wickham H, Sievert C. ggplot2 : elegant graphics for data analysis. Cham, Switzerland: Springer; 2016.

21. Khalifa KA, Badawy HM, Radwan WM, Shehata MA, Bassuoni MA. CD14(+) HLA- DR low/(-) monocytes as indicator of disease aggressiveness in B-cell non-Hodgkin lymphoma. Int J Lab Hematol. 2014;36(6):650–5.

22. Mair F, Hartmann FJ, Mrdjen D, Tosevski V, Krieg C, Becher B. The end of gating? An introduction to automated analysis of high dimensional cytometry data. Eur J Immunol. 2016;46(1):34–43.

23. Li H, Shaham U, Stanton KP, Yao Y, Montgomery RR, Kluger Y. Gating mass cytometry data by deep learning. Bioinformatics. 2017;33(21):3423–30.

24. Lux M, Brinkman RR, Chauve C, Laing A, Lorenc A, Abeler-Dorner L, et al. flowLearn: fast and precise identification and quality checking of cell populations in flow cytometry. Bioinformatics. 2018;34(13):2245–53.

25. Veglia F, Sanseviero E, Gabrilovich DI. Myeloid-derived suppressor cells in the era of increasing myeloid cell diversity. Nat Rev Immunol. 2021;21(8):485–98.

26. Lin Y, Gustafson MP, Bulur PA, Gastineau DA, Witzig TE, Dietz AB. Immunosuppressive CD14+HLA-DR(low)/- monocytes in B-cell non-Hodgkin lymphoma. Blood. 2011;117(3):872–81.

27. Azzaoui I, Uhel F, Rossille D, Pangault C, Dulong J, Le Priol J, et al. T-cell defect in diffuse large B-cell lymphomas involves expansion of myeloid-derived suppressor cells. Blood. 2016;128(8):1081–92.

28. Tadmor T, Fell R, Polliack A, Attias D. Absolute monocytosis at diagnosis correlates with survival in diffuse large B-cell lymphoma-possible link with monocytic myeloid-derived suppressor cells. Hematol Oncol. 2013;31(2):65–71.

29. Chow A, Perica K, Klebanoff CA, Wolchok JD. Clinical implications of T cell exhaustion for cancer immunotherapy. Nat Rev Clin Oncol. 2022;19(12):775–90.

30. Verma V, Shrimali RK, Ahmad S, Dai W, Wang H, Lu S, et al. PD-1 blockade in subprimed CD8 cells induces dysfunctional PD-1(+)CD38(hi) cells and anti-PD-1 resistance. Nat Immunol. 2019;20(9):1231–43.

31. Bjorkstrom NK, Riese P, Heuts F, Andersson S, Fauriat C, Ivarsson MA, et al. Expression patterns of NKG2A, KIR, and CD57 define a process of CD56dim NK-cell differentiation uncoupled from NK-cell education. Blood. 2010;116(19):3853–64.

32. Plonquet A, Haioun C, Jais JP, Debard AL, Salles G, Bene MC, et al. Peripheral blood natural killer cell count is associated with clinical outcome in patients with aaIPI 2-3 diffuse large B-cell lymphoma. Ann Oncol. 2007;18(7):1209–15.

33. Klanova M, Oestergaard MZ, Trneny M, Hiddemann W, Marcus R, Sehn LH, et al. Prognostic Impact of Natural Killer Cell Count in Follicular Lymphoma and Diffuse Large B- cell Lymphoma Patients Treated with Immunochemotherapy. Clin Cancer Res. 2019;25(15):4634–43.

